# Is competition for cellular resources a driver of complex trait heritability?

**DOI:** 10.1101/2022.03.25.485854

**Authors:** Olivier Naret, Yuval Simons, Jacques Fellay, Jonathan K Pritchard

## Abstract

Most human complex traits are enormously polygenic, with thousands of contributing variants with small effects, spread across much of the genome. These observations raise questions about why so many variants–and so many genes–impact any given phenotype. Here we consider a possible model in which variant effects are due to competition among genes for pools of shared intracellular resources such as RNA polymerases. To this end, we describe a simple theoretical model of resource competition for polymerases during transcription. We show that as long as a gene uses only a small fraction of the overall supply of polymerases, competition with other genes for this supply will only have a negligible effect on variation in the gene’s expression. In particular, although resource competition increases the proportion of heritability explained by trans-eQTLs, this effect is far too small to account for the roughly 70% of expression heritability thought to be due to trans-regulation. Similarly, we find that competition will only have an appreciable effect on complex traits under very limited conditions: that core genes collectively use a large fraction of the cellular pool of polymerases and their overall expression level is strongly correlated (or anti-correlated) with trait values. Our qualitative results should hold for a wide family of models relating to cellular resource limitations. We conclude that, for most traits, resource competition is not a major source of complex trait heritability.

## 1 Introduction

Since the advent of genome-wide association studies some 15 years ago, there has been huge progress toward determining the genetic basis of many human complex traits (Klein et al. 2005; Wellcome Trust Case Control Consortium 2007; Claussnitzer et al. 2020). However, early studies found something perplexing: namely that the lead variants for any given trait typically explained only a small fraction of the heritability that had been predicted by family studies (Weedon et al. 2008; Manolio et al. 2009). This gap between the heritability accounted for by top variants, and the heritability observed in family studies, caused so much consternation that in 2008 it was referred to as the “mystery of missing heritability” (Maher 2008).

This mystery was largely resolved when it was shown that most of the trait heritability comes from large numbers of common variants with very small effect sizes, whose signals fall far below genome-wide significance (International Schizophrenia Consortium, Purcell, Wray, Stone, Visscher, O’Donovan, Sullivan, and Sklar 2009; Yang et al. 2010). Further work since then has shown that for many complex traits, there are on the order of 10^4^ or even 10^5^ variants across the genome that affect trait variance (Zhang, Qi, Park, and Chatterjee 2018; Frei et al. 2019; O’Connor, Schoech, Hormozdiari, Gazal, Patterson, and Price 2019; Sinnott-Armstrong, Naqvi, Rivas, and Jonathan K Pritchard 2021). Although there is some contribution from coding variants, most of the heritability comes from non-coding variants impacting gene regulation (Trynka, Sandor, Han, Xu, Stranger, X. S. Liu, and Raychaudhuri 2012; Pickrell 2014; Finucane et al. 2015). These variants are spread surprisingly uniformly across the genome rather than being strongly concentrated near important genes or in particular chromosomal regions (Loh et al. 2015; Shi, Kichaev, and Pasaniuc 2016). Indeed the overall genetic architecture of most complex traits bears a striking resemblance to the classic infinitesimal model of quantitative genetics, introduced in Fisher’s 1919 paper (Fisher 1919) and characterized further mathematically by Barton et al and as summarized in Turelli’s comment(Turelli 2017; Barton, Etheridge, and Véber 2017).

However, such analyses also indicate another curious feature of the data. While the strongest GWAS signals are usually enriched near trait-relevant genes, in most cases these trait-relevant genes contribute only a small fraction of the heritability (Jostins et al. 2012; Wood et al. 2014; Boyle, Li, and Jonathan K. Pritchard 2017; Zhu and Stephens 2018; Fernández-Tajes, Gaulton, Bunt, Torres, Thurner, Mahajan, Gloyn, Lage, and McCarthy 2019). The observation that the SNPs contributing heritability are spread relatively uniformly across the genome (Loh et al. 2015) implies that a large fraction of genes must be contributing to the trait variance (Boyle, Li, and Jonathan K. Pritchard 2017). For example, a recent paper from our group examined GWAS data for three molecular traits– urate, IGF-1, and testosterone–where a great deal is known about the relevant biological pathways (Sinnott-Armstrong, Naqvi, Rivas, and Jonathan K Pritchard 2021). Aside from one major effect locus for urate, that paper concluded that in aggregate the lead biological pathways for each trait only explain about 10% of the total SNP-based heritability. Instead, for all three traits, the bulk of the heritability comes from a large number of SNPs spread relatively uniformly across the genome: we estimated around 4, 000-12, 000 causal variants for the three molecular traits and 80, 000 causal variants for height. Hence, paradoxically, for typical traits, most of the heritability appears to act mainly through seemingly trait-irrelevant genes.

### 1.1 Why do so many genes affect trait variance?

Thus, the resolution of the missing-heritability question leads to a second, and more mechanistic question: *Why are complex traits so enormously polygenic, and why do so many different genes affect trait variance?*

In two recent papers, our group proposed a simple quantitative model that we referred to as the “omnigenic” model, to explain this (Boyle, Li, and Jonathan K. Pritchard 2017; X. Liu, Li, and Jonathan K. Pritchard 2019). Summarized very briefly, this model proposes that a modest fraction of all genes have direct effects on a phenotype of interest; these are referred to as “core genes”. Meanwhile, all of the other genes expressed in trait-relevant cell types are referred to as “peripheral genes”. While the peripheral genes do not exert direct effects on the trait, by definition, the expression levels of peripheral genes can have indirect effects on the trait via gene regulatory networks. Indeed, we proposed that the large majority of the heritability actually flows through indirect trans-regulatory effects from peripheral genes.

This model is currently difficult to test directly due to our limited knowledge of gene networks and core genes. However, our analysis of molecular traits strongly supports the conclusion that core genes typically contribute only small fractions of the heritability (Sinnott-Armstrong, Naqvi, Rivas, and Jonathan K Pritchard 2021). Recent work on correlations between polygenic scores for various traits and whole blood gene expression of likely core genes also supports our model for how trait variation is mediated by a small subset of core genes (Võsa, Claringbould, Westra, Bonder, Deelen, Zeng, Kirsten, Saha, Kreuzhuber, Kasela, et al. 2018).

Furthermore, we showed that there is a natural connection between our model and estimates of cis- and trans-heritability of gene expression (X. Liu, Li, and Jonathan K. Pritchard 2019). Surveying work that measures gene expression heritability in a variety of cell types and species, we estimated that typically *∼* 70% of gene expression heritability is due to trans regulation (X. Liu, Li, and Jonathan K. Pritchard 2019). Since trans-eQTLs have very small effect sizes compared to cis-eQTLs, this implies that a typical gene must be regulated by very large numbers of trans-eQTLs–most of which lie far below the detection threshold for current studies. Based on the 70% estimate for trans heritability of expression, our model implies that peripheral genes can be expected to contribute between *∼* 70% to nearly 100% of the heritability for any given trait, depending on the number of core genes and their relative positions within the regulatory network.

It’s worth noting that other types of effects also contribute to the observed architectures of complex traits but do not resolve the paradox of extreme polygenicity, and will not be considered in detail in this paper. First, many disease endpoints are impacted by multiple separate intermediate processes, each of which is, itself, polygenic. For example, diabetes risk is affected by adiposity, lipid levels and distribution, and liver function, each of which has a polygenic basis (Udler 2019). Thus, any variants that affect the intermediate processes can potentially be detected in GWAS of the endpoint trait (Turkheimer 2000; Gottesman and Gould 2003; Pickrell, Berisa, J. Z. Liu, Ségurel, Tung, and Hinds 2016; Udler 2019). While this hierarchical nature of traits certainly contributes to the high polygenicity of some disease endpoints, it seems unlikely to be a complete and general explanation given that virtually all complex traits show high polygenicity. To give just one example, urate, which is controlled mainly by solute channels in the kidneys was estimated to have *∼* 12, 000 causal variants (Sinnott-Armstrong, Naqvi, Rivas, and Jonathan K Pritchard 2021). A second relevant effect is that selective constraint can play a “flattening” role on signals by lowering the allele frequencies of the large-effect variants (O’Connor, Schoech, Hormozdiari, Gazal, Patterson, and Price 2019). This phenomenon likely helps to explain the typically modest contributions of core genes. But at the same time we lack a mechanistic explanation for how it is that so many genes can have nonzero effects. Alternatively, non-biochemical mechanisms can also be a hidden effective trans-acting factor on gene expression. For example, a recent study using computer simulations and a polymer model of chromosomes showed that spatial correlations arising from 3D genome organization lead to stochastic and bursty transcription as well as complex small-world regulatory networks.(Brackley, Gilbert, Michieletto, Papantonis, Pereira, Cook, and Marenduzzo 2021).

### 1.2 The role of resource competition in trans-regulation and heritability

In this paper, we consider the role of a mechanism for trans regulation that is distinct from the network-based model that we considered previously (Boyle, Li, and Jonathan K. Pritchard 2017; X. Liu, Li, and Jonathan K. Pritchard 2019). In the original phrasing of our model, we assumed implicitly that the effects from peripheral genes are transmitted via specific regulatory interactions in cellular regulatory networks. Examples of specific regulatory interactions include repressors and transcription factors regulating their target genes and protein, but any type of molecular interaction between genes that affects their expression would fit within that framework (Võsa, Claringbould, Westra, Bonder, Deelen, Zeng, Kirsten, Saha, Kreuzhuber, Yazar, et al. 2021; Freimer, Shaked, Naqvi, Sinnott-Armstrong, Kathiria, Chen, Cortez, Greenleaf, Jonathan K. Pritchard, and Marson 2021). In the present paper, we consider a non-specific form of regulation: resource competition.

The foundational premise of the resource competition model is that each cell possesses limited pools of shared molecules crucial for gene expression and regulation. These molecules include RNA Polymerase II, nucleotides, spliceosomes, tRNAs, and ribosomes (Brendler, Godefroy-Colburn, Yu, and Thach 1981; Godefroy-Colburn and Thach 1981; Chu, Barnes, and Haar 2011; Walden, Godefroy-Colburn, and Thach 1981; Brendler, Godefroy-Colburn, Carlill, and Thach 1981; De Vos, Bruggeman, Westerhoff, and Bakker 2011). Recent work underscores competition as the mechanism ensuring that transcription remains consistent irrespective of the genome copy number throughout the cell cycle(Lin and Amir 2018). If an individual carries the high-expressing genotype then we can expect this to very slightly reduce the number of RNA polymerases and other shared resources available to all other genes. Hence, the existence of resource limitations implies that every cis-eQTL must act as a weak trans-eQTL for every other gene.

We should clearly expect the net effect of any single cis-eQTL to be tiny, but what about in aggregate? We know that a large fraction of genes have cis-eQTLs (GTEx Consortium et al. 2018). If there are *10*^*4*^ eQTLs in a cell type of interest, then **could these in aggregate drive a meaningful effect on the variance of any given gene, or on the heritability of a trait controlled by that cell type through resource competition?**

In this paper we use a mathematical model to show that the aggregate effects of resource competition are likely to be negligible in practice. Instead, it is more likely that most trans-acting effects on heritability flow through specific molecular interactions in gene regulatory networks.

## 2 A model for intracellular resource competition

We study these questions using a simple linear model of resource competition in a scenario of complete resource limitation: i.e., where there is a fixed resource pool, and all resources are in full use at all times. When resource competition is partial, when the limited resource is not in full use all of the time, resource competition would have an even smaller effect. To make the model specific, we describe it in terms of competition for RNA Pol II, but competition for other types of resources would be modeled very similarly.

We first examine the effect of the resource competition on the variation in expression level of a single gene and show that it can only account for a tiny fraction of the trans-regulation of gene expression. We then apply this model to a complex trait under the core gene (omnigenic) model of Liu *et al*. (X. Liu, Li, and Jonathan K. Pritchard 2019). We show that, under most plausible conditions, resource competition has only a negligible effect on the proportion of trans-heritability and on the overall trait variance.

### 2.1 A basic model of competition for polymerases

To focus on the parameters that are directly relevant here, we use a simple model that relates the expression level of a particular gene to the proportion of polymerase bound to and transcribing that gene. Thus, we absorb all the complexities of transcriptional regulation into a single parameter per gene.

We treat the binding of polymerases to promoters as multi-substrate Michaelis-Menten kinetics (Michaelis, Menten, et al. 1913; Schäuble, Stavrum, Puntervoll, Schuster, and Heiland 2013). To emphasize the connection with Michaelis-Menten, we follow the standard notation from chemistry, denoting the number of Pol II molecules bound to gene *i* (either in the promoter or gene body) per cell as the *concentration* of bound polymerase [*PG*_*i*_].

As we show in the supplement, the concentration of polymerase bound to gene *i* is

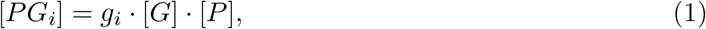

where [*G*] is the concentration of promoters (assumed constant, i.e., two copies per cell for each gene), [*P*] is the concentration of free polymerase, and *g*_*i*_ is the reciprocal of the Michaelis constant. Note that *g*_*i*_ measures gene *i*’s ability to bind free polymerase and can increase if the polymerase affinity to the gene’s promoter increases or the rate of transcription initiation increases. In the genetic context, variants that affect gene expression in cis (i.e., cis eQTLs), would do so by changing *g*_*i*_.

We denote the overall concentration of polymerase as [*P*]_0_, and assume that [*P*]_0_ is fixed. Then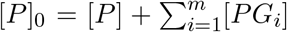 i.e., the overall concentration of polymerase is the sum of the free polymerase, [*P*], and the polymerase bound to all genes. We are interested in the limit of strong resource competition so we assume there is very little free polymerase, i.e. [*P*] *<<* [*P*]_0_, since this is the limit in which resource competition is strongest and would have the largest effect. Therefore,

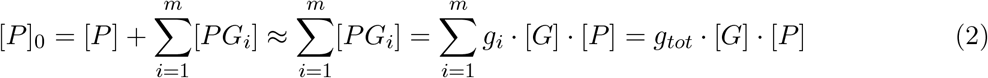

with *m* being the number of genes and 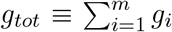 defined as a total free polymerase binding ability. Therefore,

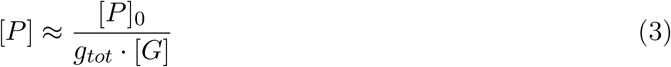

and

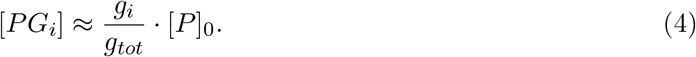

This equation shows that the proportion of time a polymerase is bound to gene *i* is proportional to 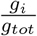 or, in other words, a fraction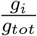 of the overall polymerase [*P*]_0_, is bound to gene *i*.

The rate of transcription of gene *i* is proportional to [*PG*_*i*_] and if we assume that the overall polymerase concentration, [*P*]_0_, is constant in time and identical between individuals then, at equilibrium, the expression level of gene *i, x*_*i*_, is

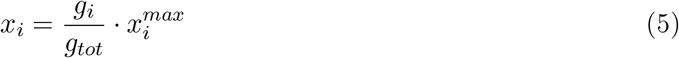

with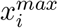 being the maximal gene expression level of gene *i* (see supplement for full derivation). 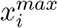 is the hypothetical expression level achieved if all the polymerases were actively transcribing gene *i*. Generally, only a small fraction of the polymerase pool is transcribing any single gene, meaning that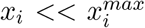. We henceforth measure each gene’s expression level in units of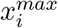, i.e., we set 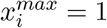 for each gene.

Under this model, gene expression of gene *i* is simply proportional to the fraction of the polymerase pool transcribing that gene. Moreover, under this scenario, Equation 5 shows that if the free polymerase binding ability *g*_*i*_ were to increase, it not only increases the number of polymerases transcribing gene *i* but also decreases the number of polymerases transcribing all other genes (by increasing *g*_*tot*_). Crucially, cis-eQTLs do exactly this: they increase (or decrease) *g*_*i*_, with small opposite-direction effects on all other genes.

**Figure 1.**
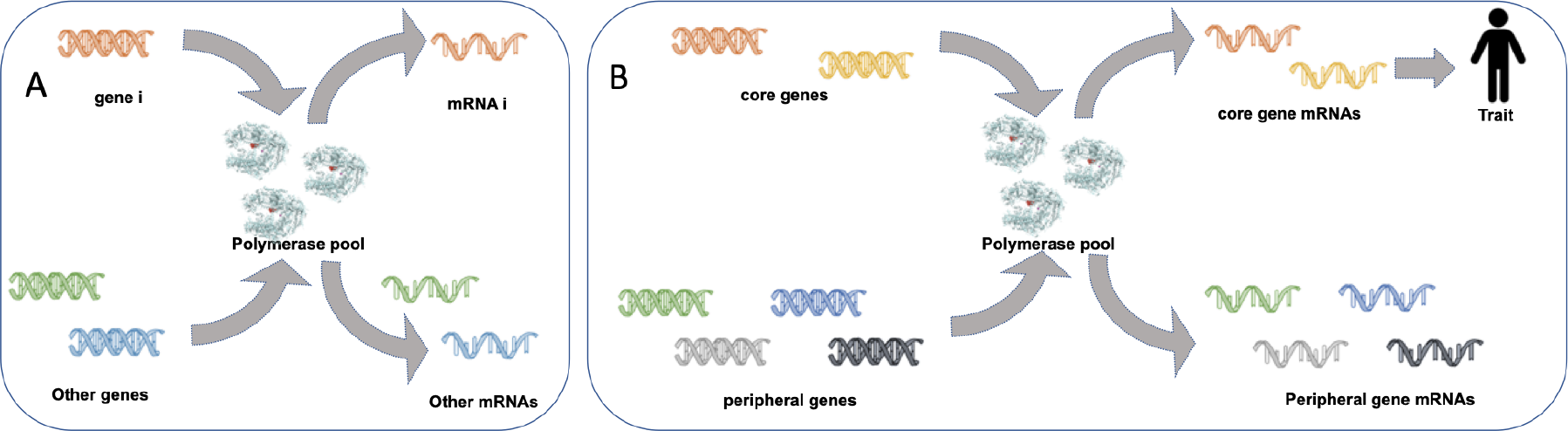
Illustration of the competition model for polymerases. **(A)** Transcription of a single gene may be affected by polymerase depletion due to transcription of other genes. **(B)** This same model extends to complex traits by distinguishing between transcription of core and peripheral genes.

Importantly, although this model is motivated by considering transcription, the functional form will hold for any form of extreme competition: gene product levels will be proportional to the gene’s share of the limited resource. For example, competition for ribosomes would lead to a similar equation for protein levels with free ribosome binding ability replacing free polymerase binding ability.

Furthermore, although we assume a relatively simple competition model for illustrative purposes, our model’s simplicity arises from assuming an extreme form of resource competition with very little free polymerase. Therefore, our key result that resource competition generally has negligible effects should be even stronger for more-complex models with less-stringent resource competition. Moreover, our results stem not from model specifics but from order-of-magnitude arguments and should therefore hold quite generally, as discussed later.

## 3 Resource competition can only explain a small fraction of trans-regulation

It is estimated that approximately 70% of the heritable variance in gene expression is due to trans-regulation (and the rest from cis-regulation) (X. Liu, Li, and Jonathan K. Pritchard 2019). Resource competition provides a possible mechanism for this, since it implies that every cis-eQTL is a (weak) trans-eQTL for all other genes. Even though for any single QTL this is a small effect, one might conjecture that trans effects could accumulate over all genes and provide a significant fraction of the estimated 70% gene expression trans heritability. However, in this section we will show that, as long as no single gene engages more than a small fraction of the overall pool of polymerases (or another limited resource), resource competition will have only a small effect on variation in gene expression and will account for only a small fraction of trans-regulation.

According to our model (5), the variance in the expression level of gene *i* across individuals is:

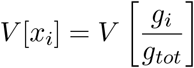

(Note that all expectations and variances henceforth are taken over individuals, i.e. over the variability in *g* values induced by genetic regulation of polymerase affinity and transcription rate.)

The variance of a ratio *V* (*a/b*), when *a << b*, can be approximated by a first-order Taylor expansion as follows (Elandt-Johnson and Johnson 1980; Kendall, Stuart, and Ord 1994):

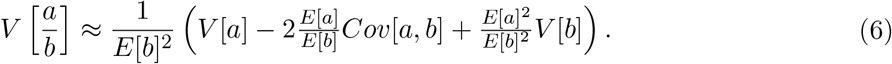

Assuming that *g*_*i*_ *<< g*_*tot*_ we can use this approximation to write:

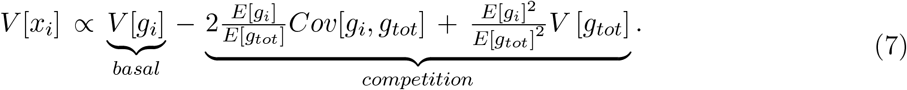

Without loss of generality, we set the proportionality constant of Equation 7 to be 1, which is equivalent to changing the units in which we measure polymerase affinity.

The first term (*V* [*g*_*i*_]) on the right side of Equation 7 reflects the various sources of genetic variance that are *not* due to resource competition: the expression variance due to cis genetic effects, environmental variance, and possibly trans effects acting via gene regulatory networks. The second and third terms reflect the effects of resource competition. Notably, the remaining two resource competition terms in this equation represent two opposite effects:

1. 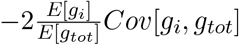 This term represents a perhaps unexpected effect. An allele associated with an increase in the free polymerase binding ability of gene *i, g*_*i*_, also leads to an increase in total free polymerase binding ability, *g*_*tot*_; hence, the impact of any cis-acting change in *g*_*i*_ is slightly counteracted by global depletion of polymerase. Thus, in this term, competition leads to a *reduction* in expression variance as represented by the second term of Equation 7.
2. 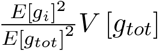 This term represents the intuitive effect that an increase (respectively, decrease) in expression of any gene soaks up some of the free polymerase, thereby slightly reducing (increasing) expression of all other genes. Thus, an increase in *V* [*g*_*tot*_] will tend to *increase* the variance of gene *i*, as represented by the third term of Equation 7.

However, critically, the magnitude of both these terms are likely to be small in practice. The two effects of resource competition depend on the ratio of the mean free polymerase binding ability of the gene of interest, *E*[*g*_*i*_], and the expected total free polymerase binding ability, *E*[*g*_*tot*_]; i.e. both effects are proportional to the fraction of polymerases bound to the gene of interest. This suggests that, as long as only a small fraction of polymerases are bound to the gene of interest, competition for polymerases would only have a small effect on variation in gene expression.

We show this explicitly using a simple but illustrative example: Assume that for all genes the mean and variance of free polymerase binding ability are identical, i.e., *E*[*g*_*i*_] = *E*_*g*_ and *V* [*g*_*i*_] = *V*_*g*_ for every *i*, and that all genes are solely under cis-regulation, i.e. *Cov*[*g*_*i*_, *g*_*j*_] = 0 if *i ≠ j*. Under these assumptions the overall variance in the expression of gene *i* is:

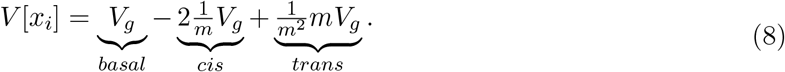

The first term in this equation is the variation due to cis-regulation of gene *i*. The second term is also a cis term and represents a dampening of cis-regulation due to the limited availability of polymerases. The third term is the trans-regulated variation in gene expression produced by resource competition. In this term, a change in the cis-regulation of any gene changes the proportion of polymerases binding to that gene and thereby mRNA production in the gene of interest.

We see that, in this simple example, resource competition leads to a reduction in the total variance but to an increase in trans regulation. However, this increase in trans regulation is inversely proportional to the number of genes, *m* ≫ 1, and is therefore tiny. This tiny effect would clearly fail to explain the roughly 70% of the variation in gene expression that is due to trans-regulation. In fact, with *∼*10, 000 genes expressed in a typical cell, the impact of resource competition would be 4 orders of magnitude smaller than the observed magnitude of trans regulation.

Relying on this simple order of magnitude argument means that the conclusion that competition has a minute effect on the overall variation and the proportion of trans-regulation is quite general. The effect of resource competition scales like 1*/m* even when genes are not identical and when there are other sources of trans-regulation. This result does not depend on which specific resource is rate limiting for transcription and holds equally for competition during translation, e.g. for ribosomes. The only possible scenarios under which competition is a large effect on gene expression variation are scenarios in which a large fraction of the pool of polymerases (or another limited resource) is bound to the gene of interest. This might plausibly happen for highly transcribed genes, such as rRNA genes; or perhaps in settings like translation that occurs in highly localized subcellular compartments with limited numbers of mRNAs.

We also conducted simulations to validate that the relative effect of resource competition is minor, on the order of 1*/m*. Details can be found in the supplementary materials, section 7.2, and are illustrated in Figures S1A and S1B.

**Figure 2.**
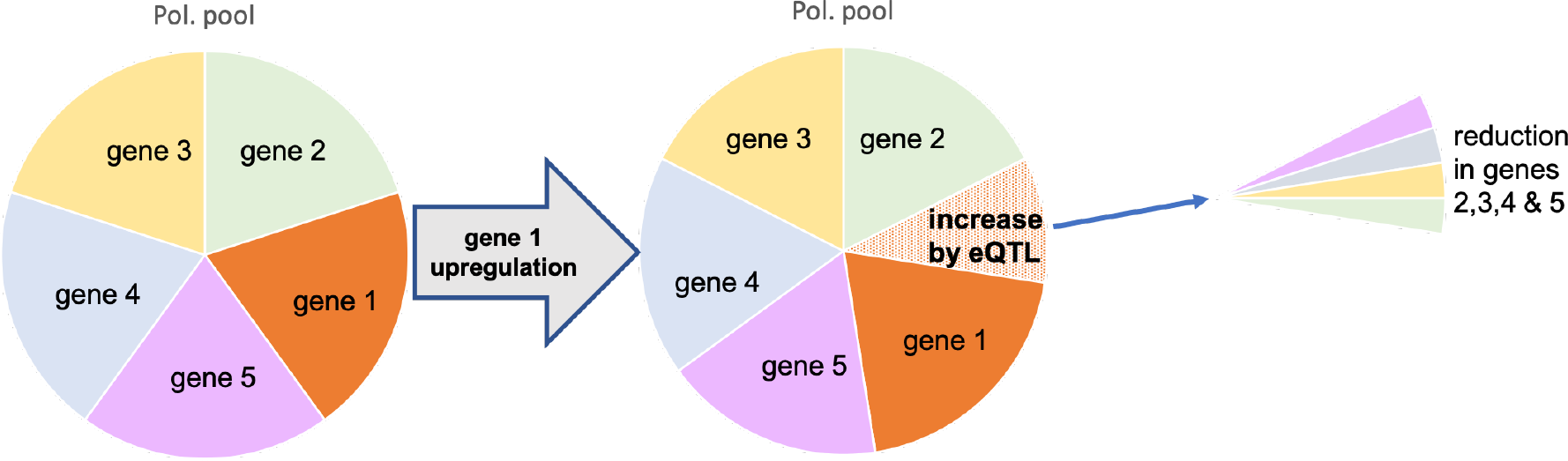
Illustration of the effect of resource competition on expression. Upregulation of Gene 1 causes a small downregulation in the expression of all other genes. Since this change is spread between all genes it scales like 1 over the number of genes.

## 4 Effect of resource competition on the variance of complex traits

Complex traits involve many genes and therefore one might hypothesize that resource competition could have a large effect on complex trait variation even though it has a small effect on the expression of any single gene. In this section, we ask **what is the impact of competition on the phenotypic variance of a complex trait?**

Using the omnigenic model laid out in Liu *et al*. (X. Liu, Li, and Jonathan K. Pritchard 2019), we consider a complex trait whose value is determined by the expression of *c* core genes. We show that there are two conditions for resource limitation to account for a significant fraction of trait variation: First, a large fraction of the pool of polymerases has to be bound to core genes. Second, an increase in the expression of core genes has to systematically increase or decrease the trait.

Following Liu *et al*., we consider a set of *c* core genes whose expression affects a complex trait, *Y*. We define the effect size per unit gene expression as *γ*, with *γ*_*i*_ being the effect size for gene *i*. A positive *γ*_*i*_ implies that an increase in the expression of gene *i* increases trait values and a negative *γ*_*i*_ indicates that an increase in the expression of gene *i* decreases trait values. In this model, an individual’s phenotype is given by:

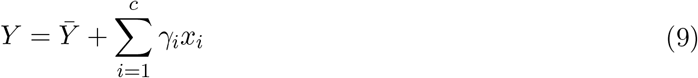

With 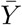being the mean phenotype in the population. We omit, for simplicity, a possible random environmental effect term.

Next, we rewrite Equation 9 in terms of free polymerase binding ability by plugging in Equation 5:

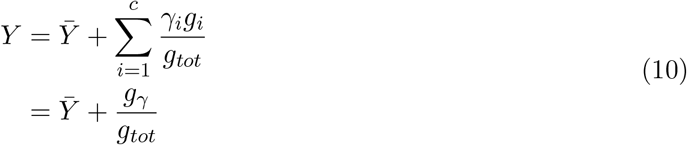

where we define, for convenience,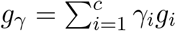.

We now use our approximation (Equation 6) to obtain an expression for the phenotypic variance:

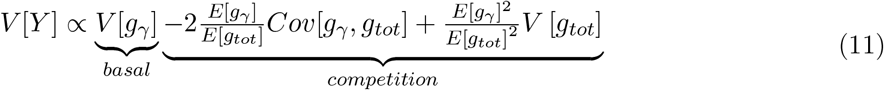

This expression is very similar in form to Equation 7 except that *g*_*γ*_ replaces *g*_*i*_. The first term now represents trait variation due to non-competitive effects, including both cis and trans regulation of core genes. The second term now represents the suppression of trait variation due to core genes competing among themselves for the limited pool of polymerases. The third term represents the increase in trait variation due to competition-induced fluctuations in the number of polymerases available for core genes.

As in the previous section (Equation 7), we can see that the effect of resource competition will depend crucially on the ratio between the averages of *g*_*γ*_ and *g*_*tot*_. This ratio is different from the ratio for the variance of a single gene seen in the previous section in two ways: (1) it concerns many core genes and not just one, and (2) each gene is associated with a *γ* value that can be either positive or negative.

To gain an intuition for the effects of these qualitative differences, we turn, once again, to the very simple model where the expression levels of all genes have identical distributions with variance *V*_*g*_ and resource competition is the only source of trans effects. Assuming that *m >>* 1 and *c >>* 1, equation 11 is then simplified to

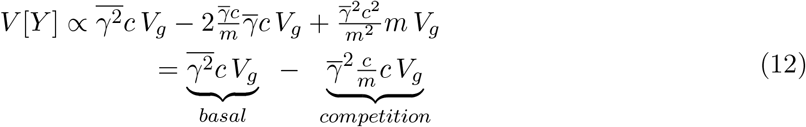

We immediately see that, in comparison to what we saw in the previous section (Equation 8), there are now two distinct reasons for the competition term to be relatively small. The effect of resource competition will be small relative to the basal term if either of the following conditions holds:

1. 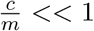, i.e. the number of core genes is much smaller than the total number of genes. As with a single gene, this is because only a small fraction of the pool of polymerases is bound to core genes, or:
2. 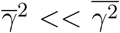, i.e. the mean effect size squared is much smaller than the mean squared effect size. This will generally be true since *γ* takes both positive and negative values. The only exception would be when trait values are strongly correlated (or strongly anti-correlated) with the overall expression level of core genes such that most *γ* values have the same sign.

This result, that resource competition is a small effect for these two reasons, should hold quite generally. In particular, it should hold if genes vary in their expression patterns, so long as core genes have comparable expression levels to peripheral genes. However, the direction of the small effect of resource competition is sensitive to such details, since they affect the balance between the two resource competition terms in Equations 7 and 12.

This result is also valid if core genes are co-regulated. As discussed by Liu *et al*. (X. Liu, Li, and Jonathan K. Pritchard 2019), if core genes tend to be co-regulated, trans effects can dominate the variance of a trait, thus inflating *V* [*g*_*γ*_], which only involves covariances between core genes, considerably compared to *Cov*[*g*_*γ*_, *g*_*tot*_] and *V* [*g*_*tot*_]. Therefore, in such a case, the relative importance of resource competition will be even smaller.

**Figure 3.**
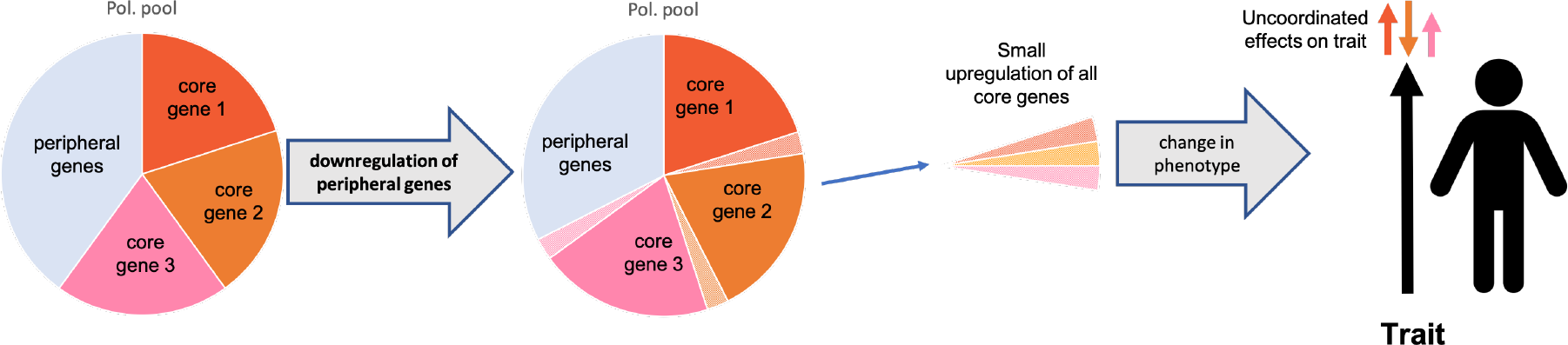
Illustration of the effect of resource competition on compex trait phenotype. Downregulation of one or more peripheral genes causes a small upregulation of all core genes. As we show in the text, this will only be an appreciable effect if core genes engage a large fraction of the overall pool of polymerases. However, as can be seen in this illustration, since an increase in core gene expression may both increase and decrease the phenotype, such an increase would only result in a minute phenotypic effect.

## 5 Discussion

In this paper, we explored the possible contribution of resource competition to gene expression and complex trait variance. Since different genes compete for the same cellular resources during transcription and translation, a variant upregulating a single gene may reduce the availability of cellular resources to all other genes. However, it is unclear, *ab initio*, if this could be a substantial effect.

We have presented a simple model of resource competition between genes, at the level of transcription. We have shown that resource competition should only have a minor effect on variation in the expression level of any given gene, as long as a small fraction of the overall pool of polymerases binds to that gene. It can therefore account for only a tiny fraction of the trans-heritability of gene expression. Similarly, only if a large fraction of the overall pool of polymerases binds to the core genes would resource competition have a major effect on trait variation. Even in such a scenario, resource competition would remain a small effect on trait variance unless trait values are strongly correlated (or anti-correlated) with the overall expression level of core genes.

While some traits may meet one of these two conditions, only a few traits should meet both. We do not know much about the expected number of core genes but it could be large for some traits. Even if a trait has a small number of core genes, these genes may bind a large fraction of the pool of polymerases in trait-relevant tissues if the expression is compartmentalized, or during specific periods of development. As for the second condition, we do not know of any category of traits for which trait values correlate with expression levels of core genes, though such traits may exist. Still, we expect that for the vast majority of traits at least one of these conditions is not met, making resource competition a negligible effect.

The work presented here is based on a very simple competition model, but one that assumes very strong competition. A more detailed model may explicitly model regulatory mechanisms, competition for multiple resources, temporal and spatial patterns, gene heterogeneity, or fluctuations. Such a model would result in a similar or weaker form of competition, with gene activity depending only partially and/or stochastically on the limited resource. Therefore, we expect our results to hold, qualitatively, even for more detailed models.

Other forms of resource competition are expected to produce near-identical results. Strong competition between genes should give the same equation 5, regardless of what is the limited resource. For example, if we consider competition for transcription factors all that would change is that *g*_*i*_ would parameterize free transcription factor binding. Similarly, if we consider competition at the level of translation instead of transcription, protein level would replace gene expression levels in Equation 5 and *g*_*i*_ could parameterize free ribosome binding (of all of gene *i*’s mRNAs). All such modes of competition, or a combination of them, would result in similar models. However it is worth noting that the conclusions may differ in contexts where a very small number of genes compete for a highly limited resource, such as access to a particular molecular transporter – this would change the relevant number of genes, *m*, such that variation in any single gene could in fact have an appreciable effect.

In summary, we have explored here resource competition as a possible contributing mechanism to expression and complex trait variation. We have laid out a foundation for understanding the effects of resource competition on the architecture of expression, protein, and trait-level variance. Our model suggests that, for most traits, competition will not be a meaningful contributor to phenotypic variance.

## 6 Acknowledgements

This work was supported by NIH grants HG011432 and HG008140 to JKP, HG011202 to YS, and by an SNSF Doc.Mobility fellowship to ON. We thank Matthew Aguirre, Hakhamanesh Mostafavi, Roshni Patel, and the entire Pritchard lab, for helpful conversations. The project was prompted in part by helpful conversations with David Botstein, Guy Sella, and Gavin Sherlock. Much of the work was conducted during an extended visit by ON to the Pritchard lab at Stanford in 2019. We would also like to thank the anonymous reviewers for their time and feedback.

## 7 Supplement: Multi-substrate Michaelis-Menten

### 7.1 Multi-substrate Michaelis-Menten

We can think of the kinetics of polymerase binding to multiple genes as following multi-substrate Michaelis-Menten kinetics, with the polymerase acting as an enzyme and the different genes as competing substrates. We present here a quick review of such kinetics, following the procedures outlined in Chou and Talalay (Chou and Talaly 1977) and arriving at a similar result to Schauble et al. (Schäuble, Stavrum, Puntervoll, Schuster, and Heiland 2013). We emphasize the assumptions we use and the simplifications arising from them.

We model mRNA production at each gene as an irreversible reaction facilitated by the polymerase, i.e.

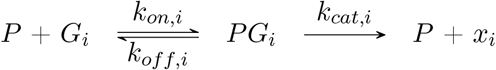

which implies that

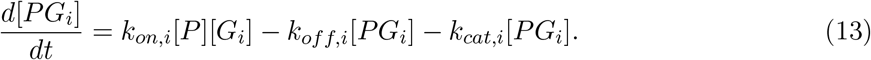

and therefore, at equilibrium, when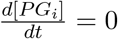,

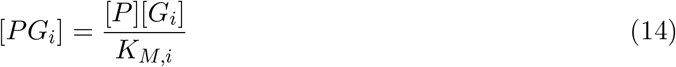

with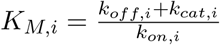being the Michaelis constant.

The overall level of the polymerase, [*P*]_0_, is constant (or externally set). Therefore,

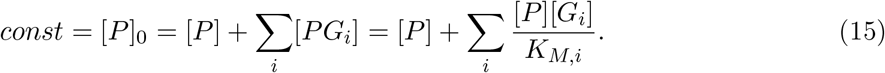

Here comes our first major assumption and simplification - since each gene has exactly two copies per cell then [*G*_*i*_] is the same for all genes, and we set [*G*_*i*_] = [*G*]. Therefore,

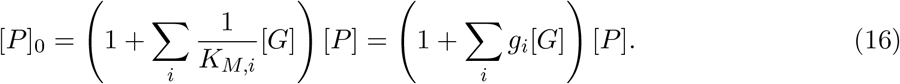

where, for convenience, we define 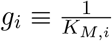.

We are interested in scenarios of resource competition, when the proportion of free polymerase is very small. That is, when polymerases spend most of their time bound to and transcribing some gene. Mathematically, this is the limit when [*P*] << ∑_*i*_[*PG*_*i*_] or

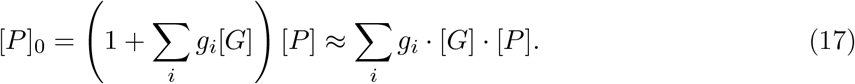

From this equation, we arrive at the results that

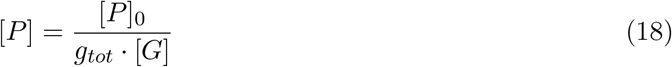

with *g*_*tot*_ = Σ_*i*_ *g*_*i*_ and therefore

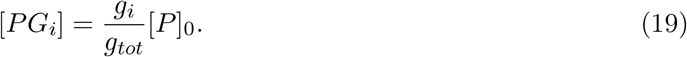

The rate of mRNA production of gene *i* is therefore

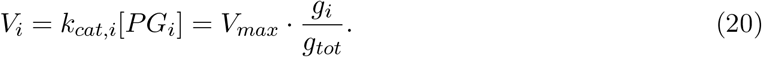

with *V*_*max*_ = *k*_*cat,i*_ *·* [*P*]_0_ being the maximum rate of mRNA production. Lastly, the level of mRNA would be

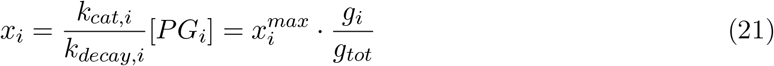

with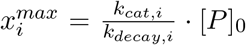 being the maximal gene expression level for this concentration of polymerase, [*P*]_0_.

### 7.2 Simulations

We simulated a dataset of 10,000 under our model under two scenarios: First, we simulated *m* identical genes with polymerase binding activity (*g*) sampled from a normal distribution with mean *μ*_*g*_ = 1000 and variance 1, and looked at variation in expression of a focal gene, marked gene *i*. In figure S1A you can see that, as predicted by equation 8, resource competition reduces the variance, and that this reduction is proportion to relative abundance of gene *i*. In the second scenario, while for all other genes *g* had a variance of 1, we varied the variance of *g*_*i*_, marked as *V*_*gi*_. As you can see in S1B, the change in expression variance is still linear in the relative abundance of gene *i* (at least for low relative abundance) but the slope depends on *V*_*gi*_. This is because changing *V*_*gi*_ changes the relative impact of the two competition terms in equation 7.

**Figure S1:**
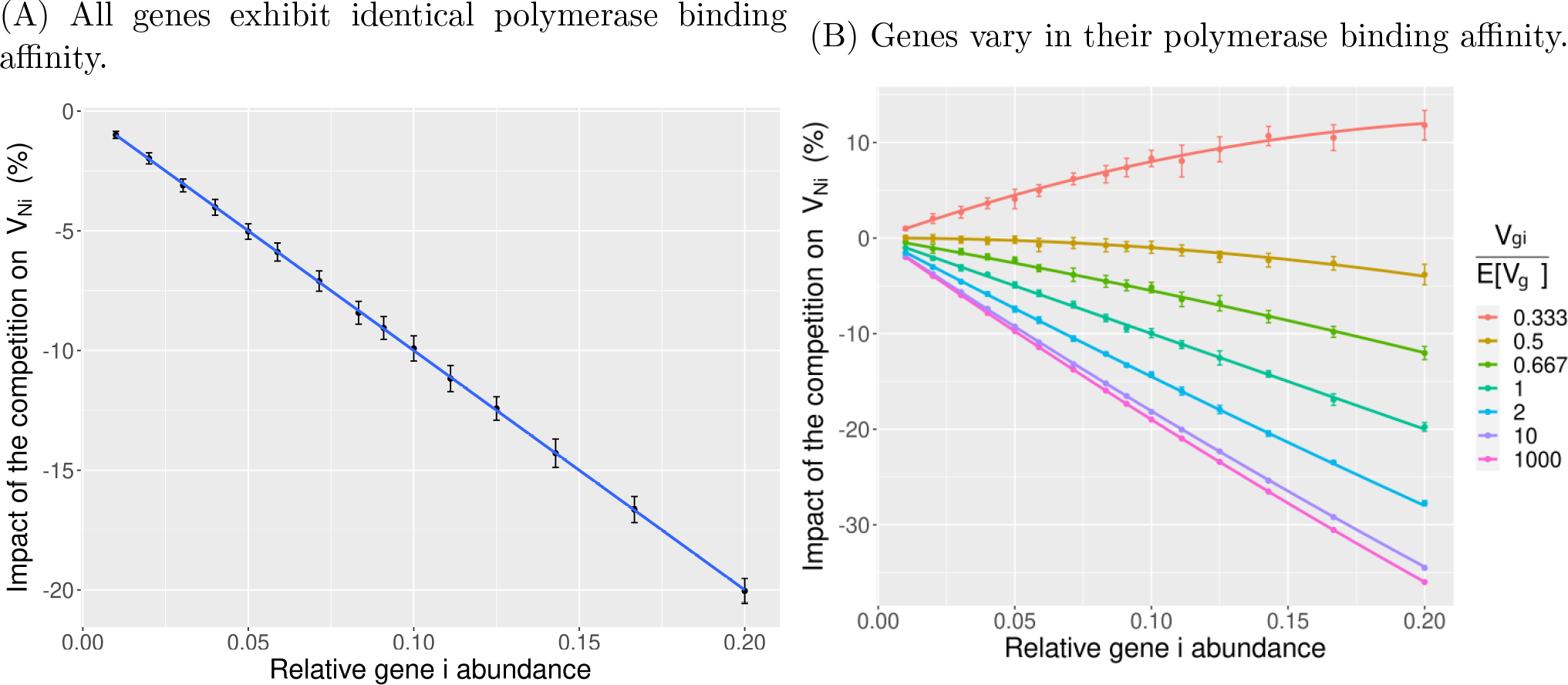
Impact of competition on variance in gene’s polymerase binding activity. **(A)** Scenario with genes having identical polymerase binding affinity across variable gene numbers *m* ranging from 5 to 100. **(B)** Scenario where genes exhibit heterogeneous polymerase binding affinities, also spanning variable gene numbers *m* from 5 to 100. The simulation data (dots with error bars) are juxtaposed with the theoretical formula (line). Each point represents the average outcome of 100 simulations based on 10,000 samples, encompassing a 90% confidence interval.

## Notes

### Competing Interest Statement

The authors have declared no competing interest.

### Summary of Updates

The manuscript is under review at eLife and we have made various minor changes to the text. The only two major differences: We added a short supplementary section with some simulations results and we have changed the order of the two co-first authors.

